# Evolutionary drivers of polymorphic sexual signals in slender anoles

**DOI:** 10.1101/2020.11.29.402784

**Authors:** Ivan Prates, Annelise B. D’Angiolella, Miguel T. Rodrigues, Paulo R. Melo-Sampaio, Kevin de Queiroz, Rayna C. Bell

**Author notes:** **Corresponding author.** Full address: Smithsonian Institution, PO Box 37012, MRC 162, Washington, DC 20013-7012. Kevin de Queiroz and Rayna C. Bell should be considered joint last (senior) author.

## Abstract

Phenotypic variation among populations, as seen in the signaling traits of many species, provides an opportunity to test whether similar factors generate shared phenotypic patterns in different parts of a species’ range. We investigate whether genetic divergence, abiotic gradients, and sympatry with closely related species explain variation in the dewlap colors of slender anoles, *Anolis fuscoauratus*. To this aim, we characterized dewlap diversity in the field, inferred population genetic structure and evolutionary relationships, assessed whether dewlap morphs are associated with climate and landscape variables, and tested for non-random associations in the distribution of *A. fuscoauratus* morphs and sympatric *Anolis* species. We found that dewlap colors vary among but not within sites in *A. fuscoauratus*. Regional genetic clusters included multiple morphs, while populations with similar dewlaps were often distantly related. Morphs did not segregate in environmental space, suggesting that dewlaps are not locally adapted to abiotic factors. Instead, we found a negative association between certain morphs and *Anolis* species with similar relative dewlap brightness, suggesting that interactions with closely related species promoted dewlap divergence among *A. fuscoauratus* populations. Slender anoles emerge as a promising system to address questions about parallel trait evolution and the contribution of signaling traits to speciation.

## Introduction

Phenotypic variation within species is ubiquitous. This variation can occur among individuals within populations, as when alternative life-history traits coexist at a given site (e.g., Sinervo and Lively 1996; Galeotti et al. 2013). Alternatively, phenotypes can vary across a species’ range. For instance, populations frequently show marked differences in coloration, vocalization, or chemical defense traits, and these phenotypes may vary across short geographic distances (e.g., Schiotz 1971; Galeotti et al. 2003; Seehausen et al. 2008; Prates et al. 2019). Some remarkable cases of polymorphism involve sexual signals, which organisms use to attract, identify, and choose suitable mates (Hill 1994; Kwiatkowski and Sullivan 2002; Jiggins et al. 2011). Variation in signaling traits within species can disrupt mate choice, and yet population differences in visual and acoustic signals are pervasive (e.g., Ryan et al. 1996; Arnqvist and Kolm 2010; Maan and Cummings 2008; Scordato and Safran 2014). Uncovering the drivers of trait variation can shed light on the ecological and evolutionary processes that underlie phenotypic diversification. Moreover, because sexual signal divergence can lead to pre-mating reproductive isolation, clarifying why signals diverge can shed light on how new evolutionary lineages arise (Jiggins et al. 2011; Hoskin et al. 2005; Gleason and Ritchie 1998).

Species with polymorphic sexual signals represent evolutionary and ecological replicates for testing hypotheses about the origins of trait diversity. One possibility is that phenotypic variation has evolved through non-adaptive processes, such as genetic drift in isolated populations (Gehara et al. 2013) or isolation-by-distance across continuously distributed populations (Campbell et al. 2010). Correspondingly, in many species, signal divergence scales directly with the time since population divergence and inversely with gene flow (e.g., Ryan et al. 1996; Bernal et al. 2005; Warwick et al. 2015). Additionally, deterministic processes, such as assortative mating, can intensify signal divergence that arises stochastically (Lande 1982; Tazzyman and Iwasa 2010). Sexual signal diversity may also be adaptive, particularly when signaling traits vary along abiotic and biotic landscape gradients. For instance, colorful signals vary with the light environment in birds and fishes (Marchetti 1993; Boughman 2001; Seehausen et al. 2008), suggesting optimized signal transmission at a local scale. Moreover, in geographic regions where two closely related lineages overlap, sexual signals may diverge via reproductive character displacement (Grant 1972). For example, certain closely related lineages of frogs produce similar calls in allopatry, but divergent calls in sympatry (Höbel and Gerhardt 2003; Hoskin et al. 2005). This pattern suggests that biotic interactions can also select for increased signal discrimination. In wide-ranging species, populations that are phenotypically similar and yet genetically divergent often occur in similar abiotic or biotic environments (reviewed in Zamudio et al. 2016). These species provide an opportunity to test whether abiotic and biotic factors lead to parallel trait evolution.

An iconic example of a diverse sexual signal is the dewlap, a colorful and extensible flap of skin positioned along the underside of the throat in *Anolis* lizards (reviewed in Losos 2009). Dewlap displays play an important role in courtship and agonistic interactions in anoles (reviewed in Tokarz 1995) and there is an extraordinary diversity of dewlap color and pattern across *Anolis* species (Nicholson et al. 2007). Some studies suggest that dewlap coloration is associated with optimizing signal transmission in relation to light environment. In the Caribbean, anole species that inhabit shaded forests more often have dewlaps with white or yellow skin color (Fleishman 1992), which reflect a high total number of photons and are thus brighter (Fleishman et al. 2009). By contrast, species from open habitats more frequently have dewlaps with red or blue skin color (Fleishman 1992), which are less reflective and, thus, darker (Fleishman et al. 2009). However, dewlap color variation might also have evolved through selection for reduced phenotypic overlap among sympatric *Anolis* species (Rand and Williams 1970; Nicholson et al. 2007), leading to character displacement. In this case, dewlap divergence may be particularly important for sympatric species that are more closely related and have more similar dorsal coloration and body sizes (Fleishman et al. 2009).

Despite the presumed role of the dewlap in species recognition and reproductive isolation, several anole species show population variation in dewlap coloration (e.g., Vanhooydonck et al. 2009; Ng and Glor 2011; Prates et al. 2015; Driessens et al. 2017; Ng et al. 2017; White et al. 2019). One species that has remarkable geographic variation in dewlap color is *Anolis fuscoauratus*, the slender anoles. In this species, males have large dewlaps with variable color phenotypes, the three most common being gray, pink, and yellow (Fig. 1). Given the wide distribution of *A. fuscoauratus* in Amazonia and the Atlantic Forest, two large disjunct rainforest blocks, this dewlap diversity might be linked to population genetic divergence and reflect incipient species. Alternatively, widespread South American anole species span pronounced environmental gradients (Prates et al. 2018), and thus dewlap morphs might be locally adapted (Ng et al. 2013a). Finally, dewlap color diversity might reflect character displacement, because *A. fuscoauratus* co-occurs with at least 15 other *Anolis* species across its distribution (Avila-Pires 1995; Ribeiro-Júnior et al. 2015; Prates et al. 2017). Therefore, the large geographic variation of dewlap color in *A. fuscoauratus* provides a promising system to test whether similar factors acting in different parts of a species’ range generate repeated phenotypic patterns.

**Fig. 1.**
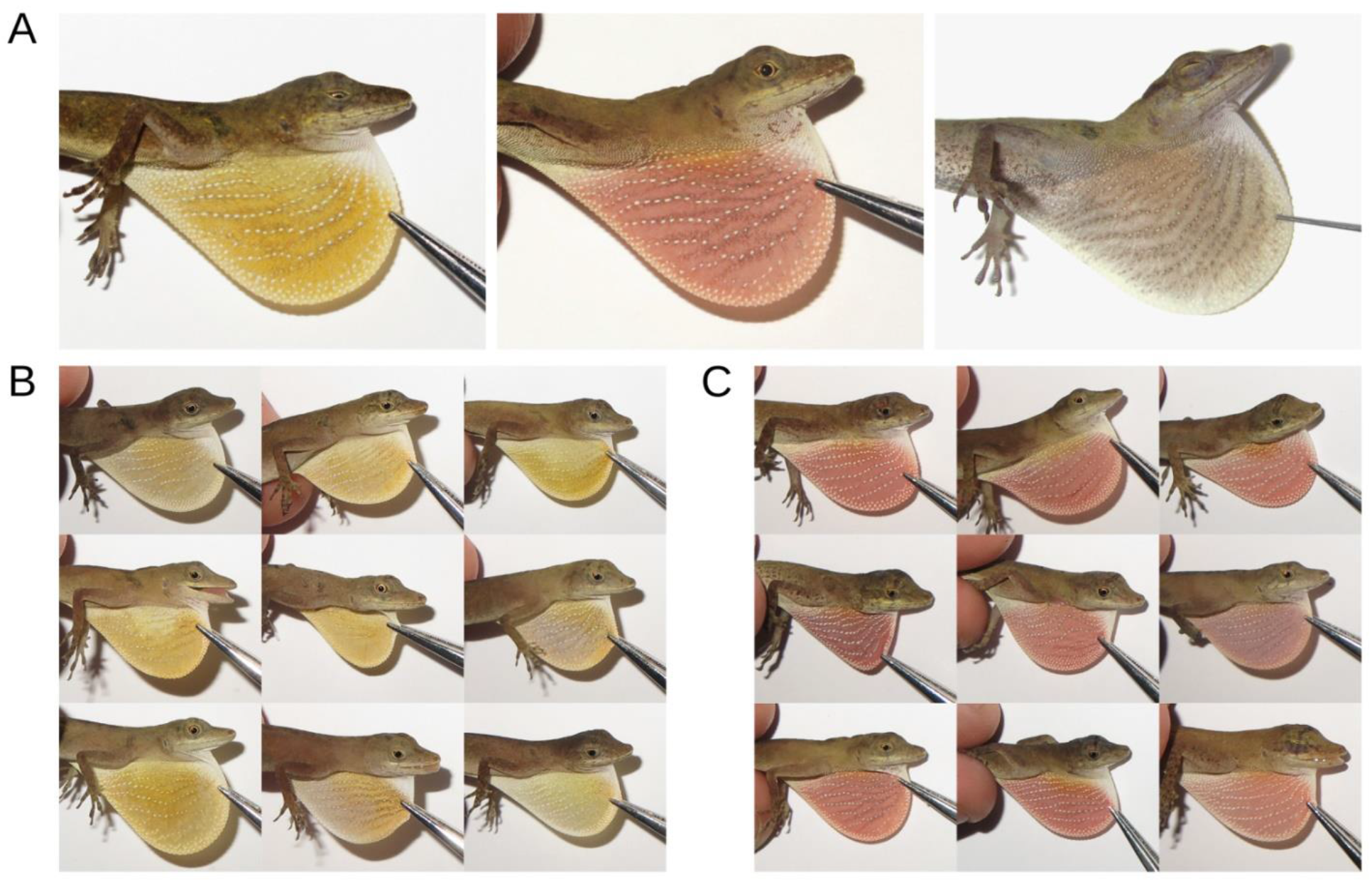
The three most common dewlap color morphs of male slender anoles, Anolis fuscoauratus: (A) yellow, pink, and gray. Example of limited intra-site dewlap color variation at (B) Rio Branco, Acre, Brazil, and (C) Senador Guiomard, Acre, Brazil; these two sites are separated by ~25 km of continuous Amazonian rainforest.

This study seeks to test whether evolutionary divergence, landscape gradients, and the composition of local *Anolis* assemblages explain the evolution of highly variable sexual signals in slender anoles. To answer this question, we recorded data on dewlap diversity and geographic variation on the basis of herpetological inventories that we performed over the last two decades in South America. We then generated genome-wide data through a reduced representation method to infer patterns of genetic structure in *A. fuscoauratus* and test whether populations with more similar dewlap phenotypes are also more closely related. To test whether dewlap coloration in slender anoles co-varies with landscape gradients over the species’ range, we estimated multidimensional environmental space occupancy by each dewlap morph based on geospatial descriptors of climate, topography, and vegetation. Lastly, we tested whether dewlap colors in this species vary as a function of local co-occurrences with other *Anolis* species, particularly those that are morphologically more similar and phylogenetically closer to *A. fuscoauratus*.

## Material and Methods

### Field assessment of dewlap variation and anole assemblage composition

*Anolis fuscoauratus* is found in both primary and secondary rainforests in South America, where it usually is the most abundant *Anolis* species locally. To characterize geographic dewlap color variation in this species, we used data from our comprehensive herpetofaunal inventories in Amazonia and the Atlantic Forest over the last two decades. No quantitative color data (e.g., spectrometric measurements) were obtained due to constraints of field sampling and infrastructure. Therefore, in our environmental and species co-occurrence analyses (see below) we only included sites for which dewlap color information were available (pictures or field notes). Based on these records, we were able to assign 32 localities to one of the three main dewlap forms most commonly seen in *A. fuscoauratus*: gray, pink, and yellow (see Results).

One of the goals of this study was to test whether dewlap variation in *A. fuscoauratus* is linked to the presence of other *Anolis* species across regions. To test this hypothesis, we obtained data on species presence at a given site based on our field inventory data. To reduce the chance of undetected species, we only included data from surveys that lasted a minimum of one week and involved at least three herpetologists searching for animals both night and day. We found all *Anolis* species expected to occur in sampled regions based on species ranges (Avila-Pires 1995; Ribeiro-Junior 2015). Attesting to the thoroughness of our sampling, over the course of our expeditions we sampled two *Anolis* species thought to be extinct and one new to science (Prates et al. 2017, 2020). The final dataset included occurrence data for other 11 anole species at the 32 sites for which *A. fuscoauratus* dewlap coloration data were available: *Anolis auratus*, *A. chrysolepis*, *A. dissimilis*, *A. nasofrontalis*, *A. ortonii*, *A. planiceps*, *A. punctatus*, *A. scypheus*, *A. tandai*, *A. trachyderma*, and *A. transversalis* (Table S1).

### Genetic sampling and data collection

For genetic analyses, individuals were collected by hand or pitfall traps, euthanized by injection of 5% lidocaine solution, fixed in 10% formalin, and preserved in 70% ethanol. Prior to fixation, a sample of liver or muscle tissue was removed and preserved in 95% ethanol. Animal handling procedures were approved by the Institutional Animal Care and Use Committee of the City University of New York and Smithsonian National Museum of Natural History. Voucher specimens were deposited in the collections of the Herpetology Laboratory and Museum of Zoology of the University of São Paulo.

To improve inferences of genetic structure and phylogenetic relationships in slender anoles, we included in the genetic analyses samples from sites with both known and unknown male dewlap color, as well as females, which do not have developed dewlaps in *A. fuscoauratus*. Our combined sampling for genetic analyses included 164 individuals of *A. fuscoauratus* sampled at 63 sites (Fig. 2A), encompassing most of the species’ distribution (Ribeiro-Júnior 2015). Most of these samples came from sites where dewlap coloration was known (N = 108). We used *Anolis auratus* (N = 2), *A. brasiliensis* (1), *A. chrysolepis* (2), *A. meridionalis* (1), *A. ortonii* (1), *A. planiceps* (2), *A. polylepis* (1), *A. quaggulus* (1), *A. scypheus* (1), and *A. trachyderma* (1) as outgroups for phylogenetic analyses based on relationships found by Poe et al. (2017). Specimen and locality information are given in Table S2.

**Fig. 2.**
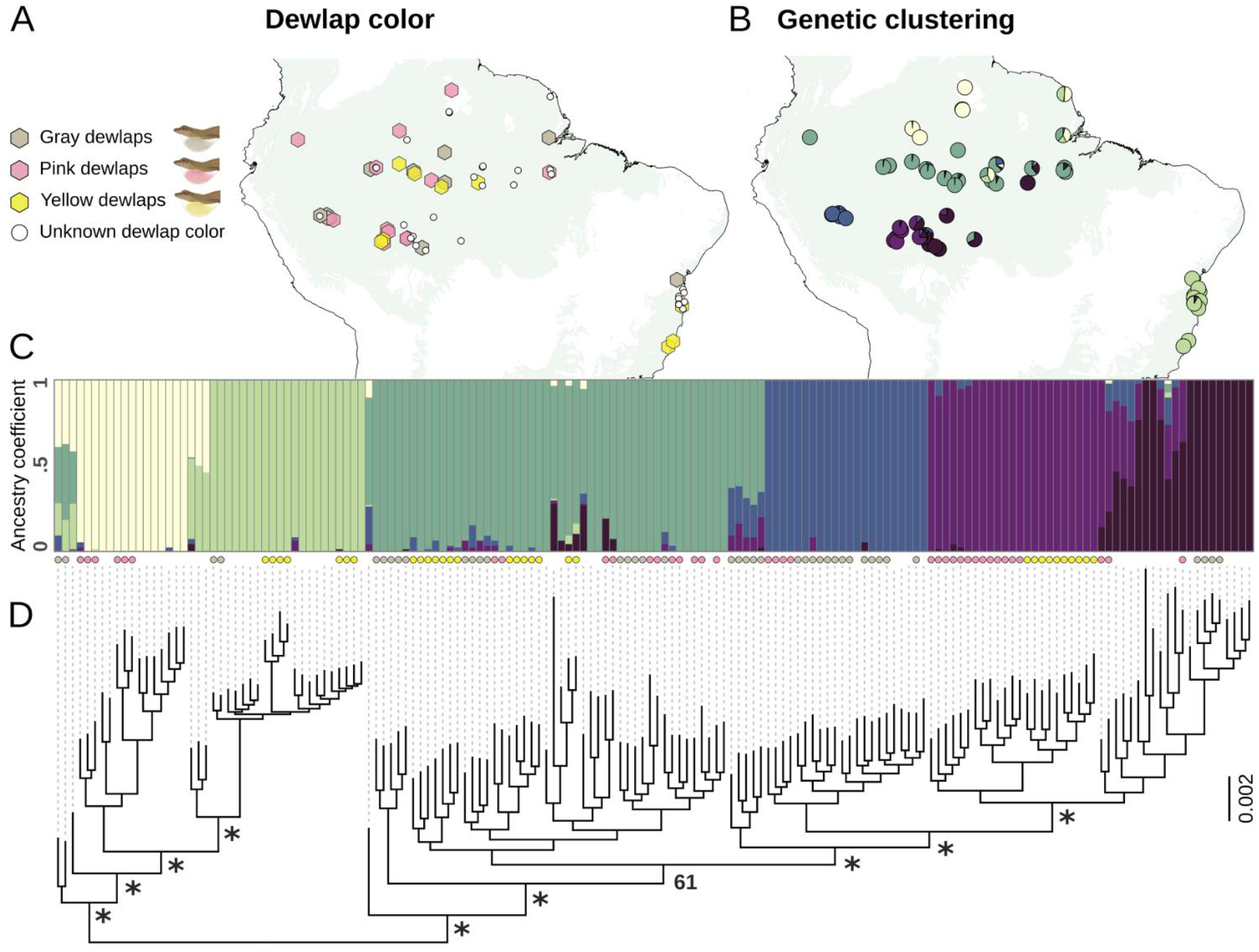
(A) Geographic dewlap color variation in Anolis fuscoauratus based on field inventories performed over the last two decades in South American rainforests. (B) Distribution of the six genetic clusters inferred from genetic clustering analysis (each cluster shown in a different color); pies indicate the average proportions of alleles (i.e., ancestry proportions) from each cluster at a given site (based on all individuals sampled at that site). Light green background on maps depicts rainforest distribution in South America. (C) Ancestry proportions from genetic clustering analyses; each bar represents an individual. When known, dewlap color at the corresponding site is indicated with colored circles below each individual bar. (D) Phylogenetic relationships among samples inferred under a Maximum Likelihood framework based on the SNP data. For clarity, nodal support is shown only for the deeper relationships between major groups inferred by genetic cluster analyses (a complete phylogeny with support of all nodes and outgroup taxa is provided in Text S2). Asterisks correspond to bootstrap support > 95.

Genomic DNA was extracted from each tissue sample through a protein precipitation extraction protocol following proteinase K and RNAase treatment (Text S1). After examining DNA fragment size using agarose gels, DNA concentration was measured using a Qubit fluorometer (Invitrogen, Waltham) and diluted to ensure a final concentration of 20–50 ng DNA per μl in a total volume of 15 μl (in TE buffer). A double-digest restriction site associated DNA library (ddRAD) (Peterson et al. 2012) was generated at the University of Wisconsin Biotechnology Center; briefly, DNA extractions were digested with the restriction enzymes PstI and MspI, and the resulting fragments were tagged with individual barcodes, PCR-amplified, multiplexed, and sequenced in a single lane on an Illumina HiSeq 2500 platform. The number of paired-end reads ranged from ~1.15 to 8.85 million per individual, with a read length of 100 base pairs. De-multiplexed raw sequence data were deposited in the Sequence Read Archive (accession [to be added at proof stage]).

### Inferring population genetic structure and evolutionary relationships

We used Ipyrad v. 0.7.30 (Eaton and Overcast 2020) to de-multiplex and assign reads to individuals based on sequence barcodes (allowing no mismatches from individual barcodes), perform *de novo* read assembly (minimum clustering similarity threshold = 0.95), align reads into loci, and call single nucleotide polymorphisms (SNPs). A minimum Phred quality score (= 33), sequence coverage (= 6x), read length (= 35 bp), and maximum proportion of heterozygous sites per locus (= 0.5) were enforced, while ensuring that variable sites had no more than two alleles (i.e., a diploid genome). Moreover, for inclusion in the final datasets, we ensured that each locus was present in at least 70% of the sampled individuals. Following the de-multiplexing step in Ipyrad, read quality and length were ensured for each sample using FastQC (available at http://www.bioinformatics.babraham.ac.uk/projects/fastqc/).

To estimate population genetic structure and admixture in *A. fuscoauratus*, we generated in Ipyrad a final dataset composed of 118,434 SNPs at 16,368 loci (including no outgroups). A single SNP was then extracted from each locus to minimize sampling of linked SNPs. We used VCFtools v. 0.1.16 (Danecek et al. 2011) to filter out SNPs whose minor allele frequency (MAF) was lower than 0.05 (Ahrens et al. 2018). After the filtering steps, 2,157 SNPs were retained across 162 individuals, with ~23 % missing data across samples. To quantify missing data, we used the Matrix Condenser tool (Medeiros and Farrell 2018). Based on the SNP data, we estimated the best-fit number of genetic clusters (K) using sNMF (Sparse Nonnegative Matrix Factorization) (Frichot et al. 2014) as implemented in the R package LEA v. 2 (Frichot and François 2015). We tested K = 1–12, with 100 replicates for each K. The run with the lowest entropy value, estimated by masking 5% of the samples, was considered to identify the best K (Frichot et al. 2014). To examine the robustness of sNMF to the regularization parameter (alpha), we ran preliminary analyses with alpha = 1, 25, 50, 100, 200, 400, 800, 1600, and 3200. Best-fit K were consistent across values of alpha, with remarkably similar model fit (entropy score range = 0.49–0.52).

To infer evolutionary relationships between inferred genetic clusters and assess whether similar dewlap colors evolved convergently in different parts of the distribution of *A. fuscoauratus*, we generated in Ipyrad a second dataset composed of 135,952 SNPs at 17,302 RAD loci (now including outgroup taxa). We performed phylogenetic inference under Maximum Likelihood on the concatenated dataset using RaxML-HPC v. 8.2.12 (Stamatakis 2014) through the CIPRES Science Gateway (Miller et al. 2010). The GTRCAT model of nucleotide evolution was used and node support was estimated with 1,000 bootstrap replicates.

### Estimating environmental space occupancy across phenotypes

Studies that performed spectral measurements *in situ* suggested that ambient light varies between forest strata (Fleishman et al. 1997; but see Fleishman et al. 2009 for a negative result). Thus, selection for dewlap detectability might vary with microhabitat use in anoles. In the case of slender anoles, previous studies found uniform microhabitat use among populations across the Amazon basin, with individuals preferably foraging on low vines, small twigs in the understory, and at the base of tree trunks (Avila-Pires 1995; Vitt el al. 2003a; Duelmann 2005). Therefore, geographic dewlap variation in *A. fuscoauratus* does not appear to be explained by differential microhabitat use among populations. We thus focus on whether sexual signal variation in this species is linked to landscape gradients at similarly large spatial scales; specifically, we test whether populations that show gray, pink, or yellow dewlaps segregate in a multidimensional environmental space defined by climate, topography, and vegetation cover. A similar approach was used to detect local adaptation in the dewlap colors of the Caribbean species *Anolis distichus* (Ng et al. 2013a).

We used 17 variables in environmental analyses (Table S3): elevation, slope, terrain roughness and terrain ruggedness (two metrics of topographic heterogeneity), mean annual cloud cover, and cover of evergreen broadleaf trees, deciduous broadleaf trees, shrubs, herbaceous vegetation, and regularly flooded vegetation (Robinson et al. 2014; Tuanmu and Jetz 2014; Wilson and Jetz 2016; Amatulli et al. 2018), all obtained from the EarthEnv database (http://www.earthenv.org); annual mean temperature, maximum temperature of the warmest month, mean temperature of the warmest quarter, annual precipitation, precipitation of the wettest month, and precipitation of the wettest quarter (Karger et al. 2017), obtained from the Chelsa database (http://chelsa-climate.org); and the climatic moisture index (a metric of relative wetness) (Title and Bemmels 2018), obtained from the ENVIREM database (http://envirem.github.io).

Values were extracted for each variable from the 32 sites for which *A. fuscoauratus* dewlap color information was available using the *Point Sampling Tool* plugin in QGIS v. 3.4.5. To visualize and compare environmental space occupancy across the range of *A. fuscoauratus*, we performed a principal component analysis (PCA) on the environmental variables and generated violin and scatter plots using R (R Core Team 2020). Moreover, we compared mean PC values between dewlap morphs based on an analysis of variance (ANOVA) using the *aov* R function. We visually inspected quantile-quantile (Q-Q) plots to detect outliers and verify that model residuals were normally distributed.

### Testing patterns of species co-occurrence

To perform a quantitative test of whether *A. fuscoauratus* dewlaps vary geographically as a function of co-occurrences with other *Anolis* species, we used the probabilistic model implemented in the *cooccur* package in R (Veech 2013; Griffith et al. 2016). To test for negative, positive, or non-significant associations between classes (e.g., species), this method calculates the observed and expected frequencies of co-occurrence between pairs of classes; the expected frequencies are calculated assuming that the distribution of a class is independent of, and random relative to, that of another class. The method returns the probabilities that lower or higher values of co-occurrence (relative to expected values) could have been obtained by chance (Griffith et al. 2016). We treated each of the three *A. fuscoauratus* color morphs and each other *Anolis* species as a distinct class.

We initially ran an analysis to test for negative co-occurrences between each of the three *A. fuscoauratus* color morphs (gray, N = 11 sites; pink, N = 12 sites; yellow, N = 9 sites) and each of the five most common co-distributed *Anolis* species. These species were detected in at least eight of the 32 sites where *A. fuscoauratus* dewlap data were available (*A. ortonii*, N = 12 sites; *A. punctatus*, N = 19 sites; *A. tandai*, N = 12 sites; *A. trachyderma*, N = 8 sites; *A. transversalis*, N = 13 sites).

We performed a second analysis grouping all 11 *Anolis* species detected in sympatry with *A. fuscoauratus* into two groups: one group (N = 24 sites) of species where males have dewlaps with bright, more reflective background skin colors (as per Fleishman 1992; Fleishman et al. 2009), namely *A. dissimilis* (white), *A. planiceps* (orange), *A. punctatus* (yellow), *A. trachyderma* (yellow and orange), and *A. transversalis* (yellow); and a second group (N = 20 sites) of species whose dewlaps have relatively darker, less reflective, high chroma background skin colors (Fleishman 1992; Fleishman et al. 2009), namely *A. auratus* (blue), *A. chrysolepis* (blue), *A. nasofrontalis* (pink), *A. ortonii* (red), *A. scypheus* (red and blue), and *A. tandai* (blue). By grouping species, we were able to incorporate data from species that occurred at fewer than eight sites. Moreover, this approach aims to accommodate the possibility that the dewlaps of *A. fuscoauratus* are influenced by multiple similar *Anolis* species jointly, rather than individual species only.

In a third analysis, we grouped the 11 sympatric *Anolis* species based not only on relative dewlap background color brightness but also on their degree of overall morphological similarity and phylogenetic relatedness to *A. fuscoauratus*. In South America, *Anolis* species belong to two major clades that diverged around 49 mya (Prates et al. 2015): the *Draconura* clade (Poe et al. 2017), a group represented in our study area by generally small, slender, brown or gray anoles, including *A. fuscoauratus* and seven other species (*A. auratus*, *A. chrysolepis*, *A. ortonii*, *A. planiceps*, *A. scypheus*, *A. tandai*, and *A. trachyderma*); and the *Dactyloa* clade (Poe et al. 2017), a group represented in our study area by four generally green or greenish-gray anoles (*A. dissimilis*, *A. nasofrontalis*, *A. punctatus*, and *A. transversalis*, the latter two attaining larger body sizes than all other sampled *Anolis*). Based on morphological similarity and evolutionary relatedness, we expect that dewlap variation in *A. fuscoauratus* might be more strongly affected by sympatric *Draconura* than *Dactyloa* species. Therefore, in this third co-occurrence analysis, we grouped *Anolis* species into three classes: *Draconura* with bright background dewlaps colors (N = 9 sites), *Draconura* with darker background dewlap colors (N = 19 sites), and *Dactyloa* (N = 23 sites).

Environmental data, species co-occurrence data, filtered genetic data, and detailed specimen information are available as Supplementary Information online and through the Dryad Digital Repository (doi: [to be added at proof stage]) and GitHub [to be deposited at github.com/ivanprates]. De-multiplexed raw sequence data were deposited in the Sequence Read Archive (accession [to be added at proof stage]). R and Unix shell scripts used to prepare and filter the data and perform all analyses are available online through GitHub.

## Results

### Dewlap variation among populations of Anolis fuscoauratus

Our field inventories found remarkable geographic variation in dewlap color over the range of *Anolis fuscoauratus*. Each of the three dewlap phenotypes (gray, pink, and yellow) was sampled in regions separated by hundreds or thousands of kilometers (Fig. 2A). Individuals from sites close to each other often had similar dewlap colors, but there were also instances of phenotypic turnover within tens of kilometers. A single color morph occurs at any given site, and intra-site variation is small (Fig. 1). Among the 32 sites with unambiguous dewlap information, anoles from 11 sites had gray dewlaps, those from 12 had pink dewlaps, and those from nine had yellow dewlaps. Three sites were visited twice over six years, at the same time of the year (January); local dewlap patterns remained the same over time. Dewlap color at a site was constant between juvenile and adult males, suggesting no ontogenetic changes.

Dewlap coloration was uniform across sites in all of the other anole species that co-occur with *A. fuscoauratus*, with the exception of two populations of *A. punctatus* (Amazon green anoles). Across most of the range of *A. punctatus*, individuals have yellow dewlaps; however, in one population from the Içá River (state of Amazonas, Brazil), the lizards had light green dewlaps, and in one from the Aripuanã River (state of Mato Grosso, Brazil) they had creamy-white dewlaps (Rodrigues et al. 2002). Of the two unique *A. punctatus* populations, only the Içá River population overlapped with the known *A. fuscoauratus* dewlaps and was included in the co-occurrence analyses (both light-green and yellow being considered brighter dewlap colors).

### Patterns of genetic structure

Cluster analyses using sNMF inferred six major genetic clusters across the geographic distribution of *A. fuscoauratus* (Fig. 2B-C). The results support admixture (as indicated by the ancestry coefficients of individuals) across genetic clusters. Dewlap phenotypes (i.e., gray, pink, and yellow) did not compose distinct genetic clusters; instead, most genetic clusters were made up of anoles with two or three different dewlap color morphs, and each morph was found in several genetic clusters. The three dewlap phenotypes occurred in both admixed and non-admixed individuals.

Inferred genetic clusters segregate in geographic space (Fig. 2B-C). Samples from the coastal Atlantic Forest formed one cluster, whereas the five remaining clusters occur in different parts of Amazonia: 1) the Guiana Shield in northern South America; 2) westernmost Brazilian Amazonia; 3) southwestern Brazilian Amazonia, west of the Madeira river; 4) south-central Brazilian Amazonia, east of the Madeira river and west of the Tapajós river; and 5) central Amazonia south of the Amazon river, extending to Ecuador in the west and to the Xingu River in the east. Admixture was inferred primarily between clusters that have adjacent geographic distributions (Fig. 2C).

### Phylogenetic patterns

Similar to the genetic cluster analyses, phylogenetic analyses inferred that each of the three *A. fuscoauratus* dewlap phenotypes (i.e., gray, pink, and yellow) do not form a distinct clade. Instead, each of the three dewlap color morphs is located in multiple parts of the tree (Fig. 2D; a phylogeny including support for all nodes and outgroup taxa is provided in Text S2).

Samples from the same site shared the same dewlap coloration and grouped together. Samples from adjacent sites also tended to group together; often, these specimens had the same dewlap color. At deeper phylogenetic levels, major clades included samples having two or three different dewlap phenotypes; within each major clade, samples with the same phenotype often were not closely related (Fig. 2D).

Mirroring the results from the genetic cluster analyses, major clades corresponded to different parts of the geographic distribution of *A. fuscoauratus*. Samples from the Atlantic Forest (indicated in lighter green in Fig. 2) and Guiana Shield in northern Amazonia (cream) group together. The clade formed by these samples is sister to a clade formed by the remaining Amazonian samples. Within the latter clade, samples from westernmost Brazilian Amazonia (in western Acre; blue) group together, as do samples from southwestern Brazilian Amazonia (in eastern Acre; lighter purple) and south-central Brazilian Amazonia (in Rondônia; darker purple). Samples from central Brazilian Amazonian (darker green) comprise two primary clades that together are paraphyletic relative to the other Amazonian clades. Relationships among these major clades generally received high support (Fig. 2D).

### Environmental space occupancy

Climate, topography, and vegetation cover vary over the distribution of *A. fuscoauratus*. For instance, annual mean temperature at sampled sites ranged from 20.2 to 26.4 ^o^C, annual precipitation from 1258 to 3511 mm, elevation from 22.5 to 913 m, and cover of evergreen broadleaf trees from 3 to 100 % (raw data for all 17 environmental variables are presented in Fig. S1). After implementing PCA on the environmental data, the first three principal components explained 37, 22, and 15 % (total of 74 %) of the environmental variation across sampled sites, respectively (PCA loadings presented in Table S4). PC1 increased with higher elevation, higher topographic complexity, and lower temperature, describing a lowland to highland axis; PC2 increased with higher precipitation and cloud cover, describing a dry to wet axis; and PC3 increased with more open and deciduous vegetation, describing an axis of evergreen forest to savanna and deciduous forest.

Plots of these first three PC axes suggest that each of the three dewlap phenotypes of *A. fuscoauratus* occur at localities that together exhibit a similar range of environmental conditions, with large overlap in environmental space (Fig. 3). An ANOVA based on all sampled sites found no significant differences between phenotypes in PC1 (p = 0.755), PC2 (p = 0.204), or PC3 (p = 0.110). After eliminating four outlier sites based on the inspection of Q-Q plots, there was a statistically significant difference in PC3 (savannah and deciduous forest to evergreen forest) across morphs (p = 0.047); however, post-hoc analyses using Tukey’s test did not support significant differences between groups in pairwise comparisons (p > 0.060).

**Fig. 3.**
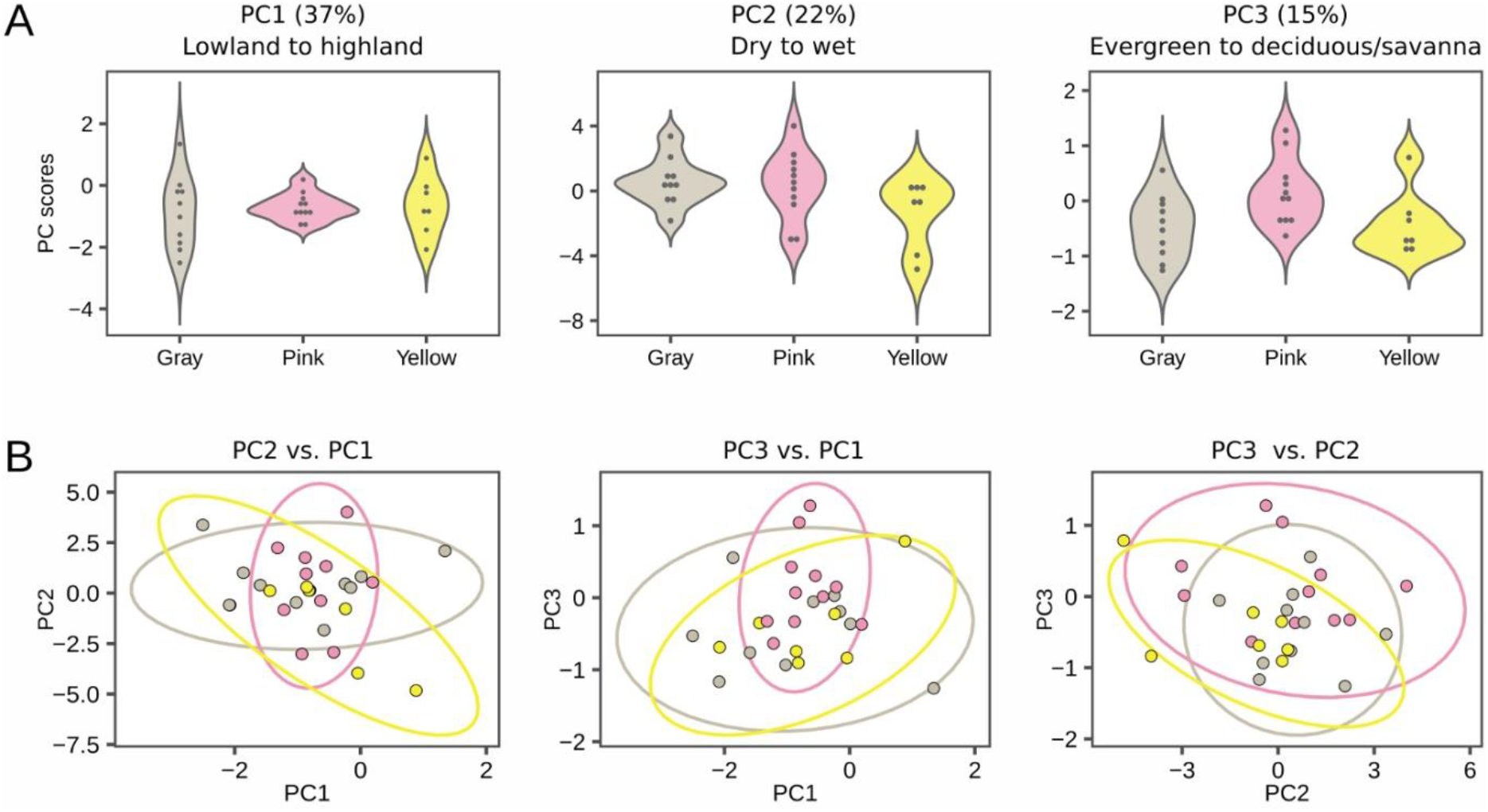
(A) Violin plots showing, for each Anolis fuscoauratus dewlap morph, the probability densities of the first three axes from an environmental PCA. PC1 describes a lowland to highland axis; PC2 describes a dry to wet axis; and PC3 describes an evergreen forest to deciduous forest and savanna axis. (B) Overlap in environmental space occupancy among dewlap morphs based on biplots of PC1, PC2, and PC3.

### Species co-occurrences

Co-occurrence analyses (Fig. 4) invariably found each of the three *A. fuscoauratus* phenotypes to be negatively associated with one another (p < 0.010), reflecting the observation of a single phenotype at each sampled site.

**Fig. 4.**
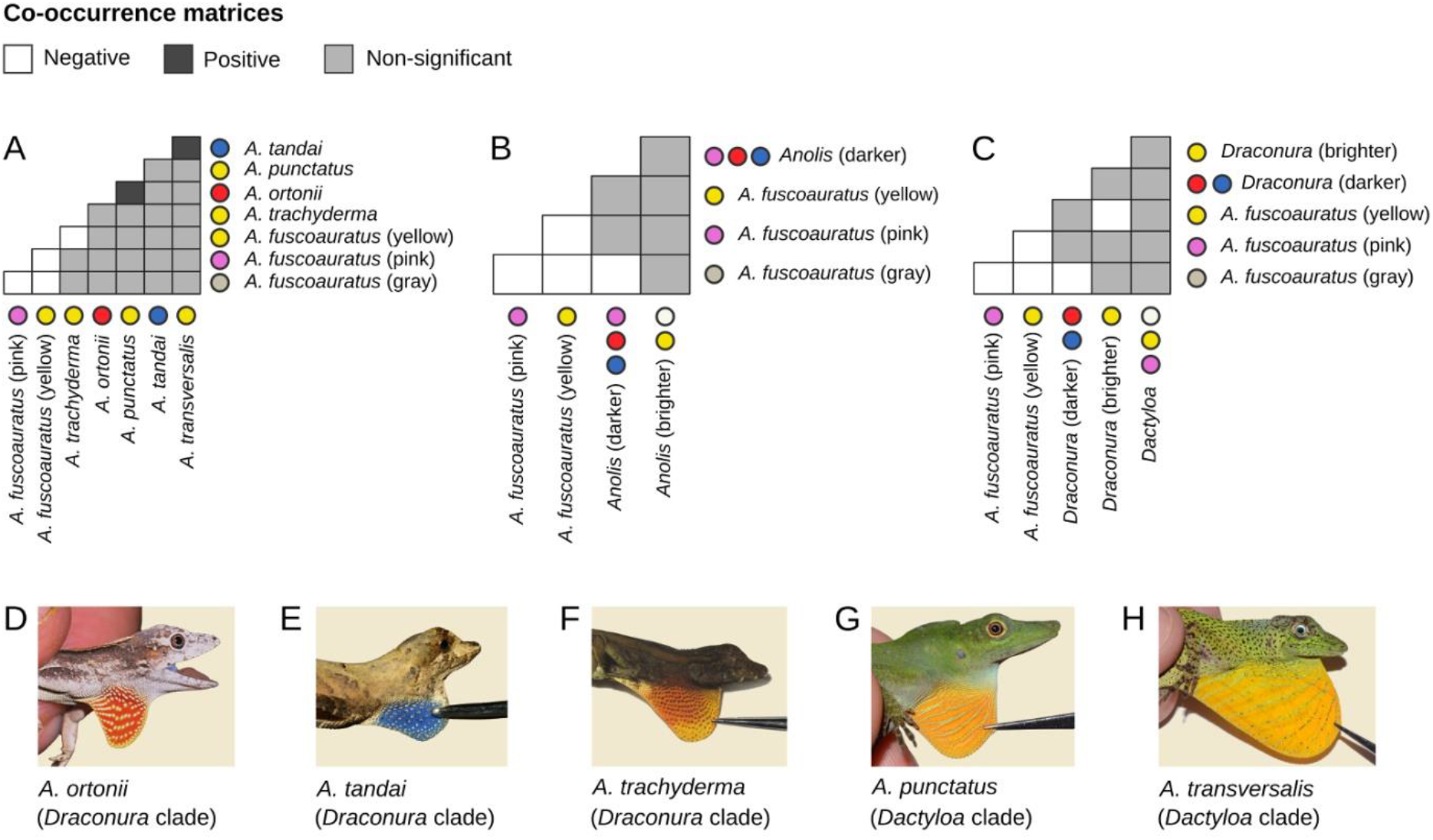
Negative, positive, or not significant patterns of co-occurrence between Anolis fuscoauratus dewlap phenotypes and sympatric Anolis species. Each square represents a pairwise comparison between each of the three A. fuscoauratus morphs (i.e., gray, pink, and yellow) and: (A) each of five co-distributed and common Anolis species (detected in at least eight out of 32 sites); (B) 11 Anolis species that occur sympatrically with A. fuscoauratus grouped into dewlaps with relatively brighter (i.e., white, yellow, yellow and orange) or darker (i.e., gray, blue, red) colors; and (C) the same 11 species grouped based on both relative dewlap color brightness and major Anolis clade (Draconura: brown or gray small, slender anoles, more similar and more closely related to A. fuscoauratus; Dactyloa: greenish larger, stockier anoles, less similar and more distantly related to A. fuscoauratus). The dewlaps of the five anole species most frequently found in sympatry with A. fuscoauratus are also illustrated: (D) Anolis ortonii, (E) Anolis tandai, and (F) Anolis trachyderma (all three in the Draconura clade); and (G) Anolis punctatus and (H) Anolis transversalis (both in the Dactyloa clade).

Analyses including the three *A. fuscoauratus* dewlap morphs and the other five most common sympatric *Anolis* species (Fig. 4A) found a negative association between *A. fuscoauratus* with yellow dewlaps and *A. trachyderma* (p = 0.047); these two classes never co-occurred. *Anolis trachyderma* and *A. fuscoauratus* both exhibit brown dorsal coloration, slender bodies, yellowish dewlaps, and belong to the *Draconura* clade of *Anolis*. The distributions of the other two *A. fuscoauratus* morphs (gray and pink) were not associated negatively (p > 0.104) or positively (p > 0.217) with the five most common sympatric *Anolis* species. This analysis also found a positive association between the occurrences of *A. tandai* (*Draconura* clade, blue dewlaps) and *A. transversalis* (*Dactyloa* clade, yellow dewlaps) (p = 0.003), as well as between *A. ortonii* (*Draconura* clade, red dewlaps) and *A. punctatus* (*Dactyloa* clade, yellow dewlaps) (p = 0.005), consistent with the observation that these species pairs frequently co-occurred at sampled sites.

When grouping the other 11 *Anolis* species based solely on relative dewlap color brightness (Fig. 4B), we found a negative association between *A. fuscoauratus* with gray dewlaps and anole species that have dewlaps with darker colors (e.g., blue and red; p = 0.005). Other relationships were not significant (p > 0.059). When grouping the other 11 *Anolis* species considering both relative color brightness and *Anolis* clade (*Draconura*: small, slender, brown anoles more similar and closely related to *A. fuscoauratus*; *Dactyloa*: greenish anoles that often attain larger body sizes and are more distantly related to *A. fuscoauratus*) (Fig. 4C), we found a negative association between *A. fuscoauratus* with gray dewlaps and *Draconura* species that also have darker dewlap colors (p = 0.011). We also found a negative association between *A. fuscoauratus* with yellow dewlaps and *Draconura* species that have brighter dewlap colors (p = 0.029). Other relationships were not significant (p > 0.071).

## Discussion

On the basis of biodiversity inventories at dozens of rainforest sites in northern South America, we found extensive dewlap color variation in *A. fuscoauratus* among sites, but limited variation within sites (Fig. 1). Similar dewlaps occur at sites hundreds or thousands of kilometers apart. In some cases, these sites are separated by unsuitable habitat (Fig. 2); for instance, yellow and gray dewlaps occur in both Amazonia and the Atlantic Forest, two rainforest regions separated by open and dry grasslands and scrublands in which slender anoles do not occur. A reduced representation genomic dataset indicated that phenotypically similar populations are often not closely related (Fig. 2), consistent with a history of repeated origin (or loss) of each of the three dewlap morphs. Moreover, a genetic cluster analysis indicated mismatches between genetic and phenotypic structure: clusters were composed of individuals with different dewlap colors, and each dewlap morph was distributed across multiple genetic clusters (Fig. 2). Estimates of environmental space occupancy found no separation by phenotype (Fig. 3), providing no support for the hypothesis of local adaptation to abiotic landscape gradients. By contrast, dewlap variation seems to be linked to the presence of other *Anolis* species across the geographic distribution of *A. fuscoauratus*. Specifically, co-occurrence analyses found that *A. fuscoauratus* with yellow (bright) and gray (darker) dewlaps occur less frequently than expected at sites where sympatric species also have brighter or darker dewlap colors, respectively (Fig. 4).

### Population isolation and sexual signal divergence

A pattern of geographically clustered phenotypic variation, as seen in slender anoles, could be generated by genetic isolation between populations due to stochastic or non-adaptive evolutionary processes. For instance, genetic drift can lead to the fixation of alternative phenotypes in isolated populations, a process that has been invoked to explain sexual signal divergence in island species (Gehara et al. 2013). This scenario predicts genetic discontinuity (i.e., allele frequency differences) between phenotypically distinct populations. However, our cluster analyses often inferred different morphs of *A. fuscoauratus* as part of the same genetic cluster, which is not consistent with a scenario of phenotypic divergence between genetically isolated populations. Alternatively, trait diversity can arise as a result of isolation-by-distance. In this case, phenotypic divergence is predicted to correlate with geographic separation (Campbell et al. 2010). However, our field surveys found phenotypic turnover among sites separated by only tens of kilometers across rainforest habitat, with no apparent geographic features that could constitute barriers to gene flow. Moreover, distant regions in distinct genetic clusters often show similar phenotypes. Taken together, these findings suggest that genetic or geographic isolation is insufficient to explain the sexual signal diversity seen in slender anoles.

### Environmental factors and sexual signal variation

A pattern of phenotypic divergence not accompanied by genetic divergence, as seen in *A. fuscoauratus*, can result from environmental factors that vary geographically. For instance, dewlap color variation might stem from local differences in diet. In birds and fishes, pigments that bestow yellow, orange, and red coloration depend on food sources of carotenoids (Endler 1983; Hill et al. 2002). In the case of anoles, these colors can also be produced via endogenously synthesized pteridins (Macedonia et al. 2000; Steffen and McGraw 2007; Alfonso et al. 2013). Experiments with *A. distichus* and *Anolis sagrei* found no change in dewlap color or pattern under alternative dietary regimes of carotenoid supplementation (Steffens et al. 2010; Ng et al. 2013b). Moreover, breeding experiments showed that dewlap coloration is heritable in *A. distichus* and *A. sagrei* (Ng et al. 2013b; Cox et al. 2017). While these studies suggest that dewlap colors are genetically determined and not plastic in *Anolis*, no such data are currently available for *A. fuscoauratus*. Future experimental studies could elucidate whether dietary pigments contribute to dewlap diversity in slender anoles, which show higher levels of geographic color variation than the previously studied species.

Alternatively, mismatches between genetic and phenotypic structure, as seen in slender anoles, may stem from adaptive divergence with gene flow (Zamudio et al. 2016). In Caribbean anoles, highly reflective (bright) dewlap colors (e.g., white and yellow) are more frequent in species that inhabit dense forests, while less reflective (darker, high chroma) dewlap colors (e.g., red and blue) appear more common in species from dry scrublands (Fleishman 1992). A similar pattern may hold for closely related populations. In *Anolis cristatellus* and *A. distichus*, dewlap spectral properties co-vary with habitat type, suggesting that signaling traits are locally adapted for increased detectability (Leal and Fleishman 2004; Ng et al. 2013a). Importantly, local adaptation in sexual signals can disrupt mate choice and promote reproductive isolation among populations, leading to speciation through sensory drive (reviewed in Boughman 2002). However, we found no association between dewlap color and spatial gradients of climate, topography, and vegetation cover in *A. fuscoauratus*. This result is inconsistent with predictions from local adaptation to landscape gradients as a driver of sexual signal diversity. It is worth noting that slender anoles are restricted to moist forests; the driest habitats where we sampled this species were forest patches on forest-savanna transitional areas (e.g., in Brazil’s state of Roraima). By contrast, the ranges of previously studied species span from mesic forests to open xeric habitats (Leal and Fleishman 2004; Ng et al. 2013a). Thus, environmental factors may be a more important driver of dewlap variation in *Anolis* species that are more ecologically diverse than *A. fuscoauratus* (Fleishman et al. 2009).

### Species co-occurrence and sexual signal divergence

Our results suggest that sexual signal variation in slender anoles may be tied to spatial turnover in the composition of ecological assemblages. Specifically, we found negative associations between the distributions of the gray and yellow *A. fuscoauratus* morphs and of *Anolis* species with similarly bright or dark dewlap colors. These associations may be influenced by the degree of morphological similarity and evolutionary relatedness among species. For instance, we found negative associations between *A. fuscoauratus* dewlap morphs with other Amazonian *Draconura* species, which also have brown or gray dorsa and slender bodies, but not with Amazonian *Dactyloa* species, which are green and stockier. These findings suggest that dewlap colors in *A. fuscoauratus* may adapt to reduce sexual signal similarity with co-distributed species at a local scale, potentially decreasing the frequency of cross-species interactions (Rand and Williams 1970). Other studies have invoked reproductive character displacement to explain diverging signaling traits among closely related lineages in sympatry, as seen in the vocalizations of birds and frogs (Wallin 1986; Höbel and Gerhardt 2003; Hoskin et al. 2005; Kirshel et al. 2009). Moreover, our results are consistent with studies showing that local selective regimes can lead to phenotypic mosaics when species interactions vary geographically (Thompson 2005; Brodie Jr et al. 2002).

Our co-occurrence results pose the question of why *A. fuscoauratus* seems to be the only Amazonian *Anolis* whose signaling traits vary as a function of the distributions of closely related species. One possibility is that dewlap diversity in slender anoles is related to behavioral relationships of dominance among species. *Anolis fuscoauratus* is the smallest and most slender of lowland Amazonian *Anolis* species (Avila-Pires 1995; Prates et al. 2017, 2020). By evolving divergent dewlaps, *A. fuscoauratus* might reduce agonistic interspecific interactions and thus avoid aggression from its larger relatives. Integrative behavioral and phenotypic analyses can be used to test the hypothesis that body size predicts dominance (or subordination) and dewlap color divergence among sympatric *Anolis* species, in Amazonia and elsewhere.

### Sexual signal divergence and reproductive isolation

Our genetic analyses suggest that patterns of genetic structure do not match phenotypic structure among populations of slender anoles, contradicting the expectation that populations with distinct sexual signals are reproductively isolated. Yet, it is widely accepted that the dewlap plays a key role in pre-mating reproductive isolation in *Anolis* lizards (reviewed by Tokarz 1995; Losos 2009). Supporting this view, behavioral experiments with *A. cybotes*, *A. marcanoi*, and *A. grahami* found stronger responses of individuals to dewlap displays of their own species than to those of other species (Losos 1985; Macedonia and Stamps 1994). Nevertheless, it is unclear whether dewlap divergence can ultimately disrupt gene flow between lineages. In the case of slender anoles, different dewlap morphs are present within each of the six genetic clusters across the species range, suggesting that genetic divergence within *A. fuscoauratus* is not associated with differences in dewlap coloration at broad or narrow spatial scales. Signal variation among interbreeding populations has been documented in other *Anolis* species (Thorpe and Stenson 2003; Stapley et al. 2011; Ng and Glor 2011; Ng et al. 2017) as well as other organisms that rely on visual signals, such as birds and fishes (Hermansen et al. 2011; Morgans et al. 2014). These studies support the idea that divergent signaling traits do not necessarily impose strong barriers to interbreeding and gene flow, even when sexual signals are locally adapted (Muñoz et al. 2013; Ng et al. 2016).

### Concluding remarks

On the basis of phenotypic, genetic, and ecological data, this study found evidence that dewlap colors in a widespread anole species are negatively associated with the local occurrence of similar and closely related species. Our finding of extensive mismatches between genetic and phenotypic structure in slender anoles at both broad and narrow spatial scales raise questions about the presumed role of the dewlap in reproductive isolation (Tokarz 1995; Losos 2009). It is worth noting that, despite dewlap color variation, populations across the range of *A. fuscoauratus* have homogeneous hemipenes, a trait linked to reproductive isolation in lizards (D’Angiolella et al. 2016). Our results also call into question the extent to which dewlap coloration is informative for species delimitation and taxonomy in *Anolis* (Prates et al. 2015).

This investigation highlights several knowledge gaps to be addressed by future studies. First, we still know little about how divergent signals affect agonistic interactions and mate choice in *Anolis*, which will require additional behavioral experimentation (e.g., Losos 1995). Moreover, the genetic basis of dewlap color variation remains unclear. Genomic analyses of phenotypically diverse species can elucidate the genetic mechanisms behind parallel trait evolution, including the contribution of standing genetic variation (reviewed in Zamudio et al. 2016) and differential gene flow across genomic regions in the face of selection (reviewed in Harrison 2012; Harrison and Larson 2014). The geographically polymorphic dewlaps of slender anoles emerge as a promising system to address these questions. Future investigations will benefit from quantitative assessments of trait variation, behavioral experiments, and comparative genomic analyses.

## Acknowledgments

We thank Anna Penna, Antoine Fouquet, Dante Pavan, Francisco Dal Vechio, João Tonini, José Cassimiro, José Mario Ghellere, Marco Sena, Mauro Teixeira Jr., Pedro Peloso, Renato Recoder, Roberta Damasceno, Sergio Marques Souza, Tuliana Bruges, Victor Prates, Vinicius Xavier, and the late Gabriel Skuk for field support. Frederick Sheldon provided samples deposited at the Louisiana State University Museum of Natural Science. Suggestions by Edward Myers, Kevin Mulder, Kyle O’Connell, Michael Yuan, and Ryan Schott greatly improved this manuscript. We thank Ana Carnaval and the members of the EEB Graduate Program at the City University of New York for discussions at the initial stages of this project. All of the laboratory and part of the computer work was conducted in and with support of the LAB facilities of the National Museum of Natural History (NMNH); we especially thank Jeffrey Hunt and Matthew Kweskin. Instituto Chico Mendes de Conservação da Biodiversidade issued collection permits (SISBIO 36753‐1, 36753‐4 and 27290‐3). This work was partially funded by FAPESP (BIOTA 2013/50297‐0), NSF (DEB 1343578), NASA through the Dimensions of Biodiversity Program, and an Associate Director of Science award at the National Museum of Natural History. IP acknowledges additional funding from a Smithsonian Peter Buck Postdoctoral Fellowship. ABD acknowledges additional funding from CNPq doctoral fellowship 142466/2011-5. MTR acknowledges additional funding from FAPESP grants 03/10335‐8 and 11/50146‐6.

## Author contributions

IP, ABD, and MTR conceptualized the study. IP, MTR, KdQ, and RCB acquired funding. IP, MTR and PRMS designed data collection and obtained the data. IP, ABD, KdQ, and RCB designed the analyses. IP wrote computer scripts, performed the analyses, prepared figures, and led the writing of the initial draft. IP, ABD, MTR, PRMS, KdQ, and RCB interpreted the results and wrote the final draft.

## Data Accessibility Statement

Environmental data, species co-occurrence data, filtered genetic data, and detailed specimen information are available as Supplementary Information online and through the Dryad Digital Repository database (doi: [to be added at proof stage]) and GitHub [to be deposited at github.com/ivanprates]. De-multiplexed raw sequence data were deposited in the Sequence Read Archive (accession [to be added at proof stage]). R and Unix shell scripts used to prepare and filter the data and perform all analyses are available online through GitHub.

## Conflict of Interest Statement

The authors declare no conflict of interest.

## Supporting Information

**Figure S1.** Violin plots depicting the ranges of all 17 environmental variables.

**Table S1.** Locality information for *Anolis fuscoauratus* and sympatric *Anolis* species used in the co-occurrence analyses.

**Table S2.** Specimen and locality information of individuals used in the genetic analyses.

**Table S3.** Locality information and data used in the environmental analyses.

**Table S4.** Loadings of variables used in environmental principal component analyses.

**Text S1.** Protein precipitation protocol used for genomic DNA extraction.

**Text S2.** Phylogenetic tree including node support values and outgroup taxa.

## Notes

### Competing Interest Statement

The authors have declared no competing interest.

