## Supplementary figures, tables, and data for "Evolutionary drivers of polymorphic sexual signals in slender anoles": Text_S1_extraction_protocol.pdf

### DNA extraction using protein precipitation for high yield DNA

**What you need:**
- Cell Lysis Buffer (step 1).
- Proteinase K (20 mg/ml) (step 1).
- RNAse A (4 mg/mL) (step 1).
- Protein Precipitation Solution (step 2).
- Ethanol (100% and 70%) (step 2).
- 1x TE or ddH2O (for elution) (step 2).
- Sterile 1.5 mL microcentrifuge tubes (two times more than the number of samples).

**1st part (cell lysis and RNAse treatment), done on: ___/___/___**
☐ Add 300 µl of Cell Lysis Solution to a 1.5 mL **labeled** tube.
☐ Add 10 µl of Proteinase K.
☐ Add 2 µl RNAse (10 ng/ul).
☐ Place the tissue sample in the tube.
☐ Incubate until dissolved at 55C (3 hours to overnight – check degree of tissue digestion).

**2nd part (protein precipitation), done on: ___/___/___**
☐ Add 100 µl Protein Precipitation Solution to the cell lysate mixture. Vortex vigorously.
☐ Centrifuge at 13000 RPM for 10min. Repeat if the pellet is not tight.
☐ Add 1000 µl of 100% EtOH to a new 1.5 mL tube.
☐ Pour off the supernatant (**=> which contains your DNA**) into this tube. Mix by inverting gently 50 times.
☐ Centrifuge at 13000 RPM for 10 min.
☐ Pour off the supernatant carefully, make sure to keep the pellet (**this is your DNA**).
☐ Add 1000 µl of 70% EtOH and invert the tube several times to wash the pellet.
☐ Centrifuge at 13000 RPM for 10 minutes.
☐ Pour off the supernatant carefully, make sure to keep the pellet (**this is your DNA**).
☐ Airdry for 10min to let all the EtOH evaporate. Some liquid might stay but that is hopefully the 30% water. Do not overdry as then the pellet will become hard to elute.
☐ Add 30 µl of 1x TE or ddH20.
☐ Incubate overnight at room temperature overnight to get all the DNA eluted. Store.

**Note:** TE is slightly better at resuspending the pellet due to its pH and is also better for long term storage. But if you will use it soon afterwards and especially if you need to dry it down its better to use ddH2O so you do not concentrate the buffer.

**Samples:**

| | | | | |
|---|---|---|---|---|
| **1** | | **13** | | **25** |
| **2** | | **14** | | **26** |
| **3** | | **15** | | **27** |
| **4** | | **16** | | **28** |
| **5** | | **17** | | **29** |
| **6** | | **18** | | **30** |
| **7** | | **19** | | **31** |
| **8** | | **20** | | **32** |
| **9** | | **21** | | **33** |
| **10** | | **22** | | **34** |
| **11** | | **23** | | **35** |
| **12** | | **24** | | **36** |

**Solutions:**
These solutions can be bought from QiaGen directly. Prices as per late 2018.

All reagents are here: https://www.qiagen.com/us/shop/lab-basics/buffers-and-reagents/puregene-accessories/#orderinginformation

**Cell Lysis Buffer:**
125 mL (Cat No. 158906) is $87 and enough for ~415 extractions (21c per sample)
1000 mL (Cat No. 158908) is $453 and enough for ~3300 extractions (14c per sample)

**Protein Precipitation Solution:**
50 mL (Cat No. 158910) is $87.40 and enough for ~500 extractions (17.5c per sample)
350 mL (Cat No. 158912) is $399 and enough for ~3500 extractions (11.4c per sample)

**Proteinase K:**
Many options not sure what is best. I used the one from the DNeasy kit.
2 mL (Cat No. 19131) is $96.30 and is enough for ~200 extractions (48c per sample)
10 mL (Cat No. 19133) is $326 and is enough for ~1000 extractions (32.6c per sample)

Total excluding ethanol/plastics is 76.5c when buying the smaller versions and 58c when buying in bulk. That compared to Qiagen DNeast kit for 250 samples which is ~700 and thus $2.8 per sample (!). Yields are also substantially higher than with the DNeasy kit.
