## Supplementary figures and images for "Evolutionary drivers of polymorphic sexual signals in slender anoles"

### Fig_S1_plot_environmental_variables.pdf

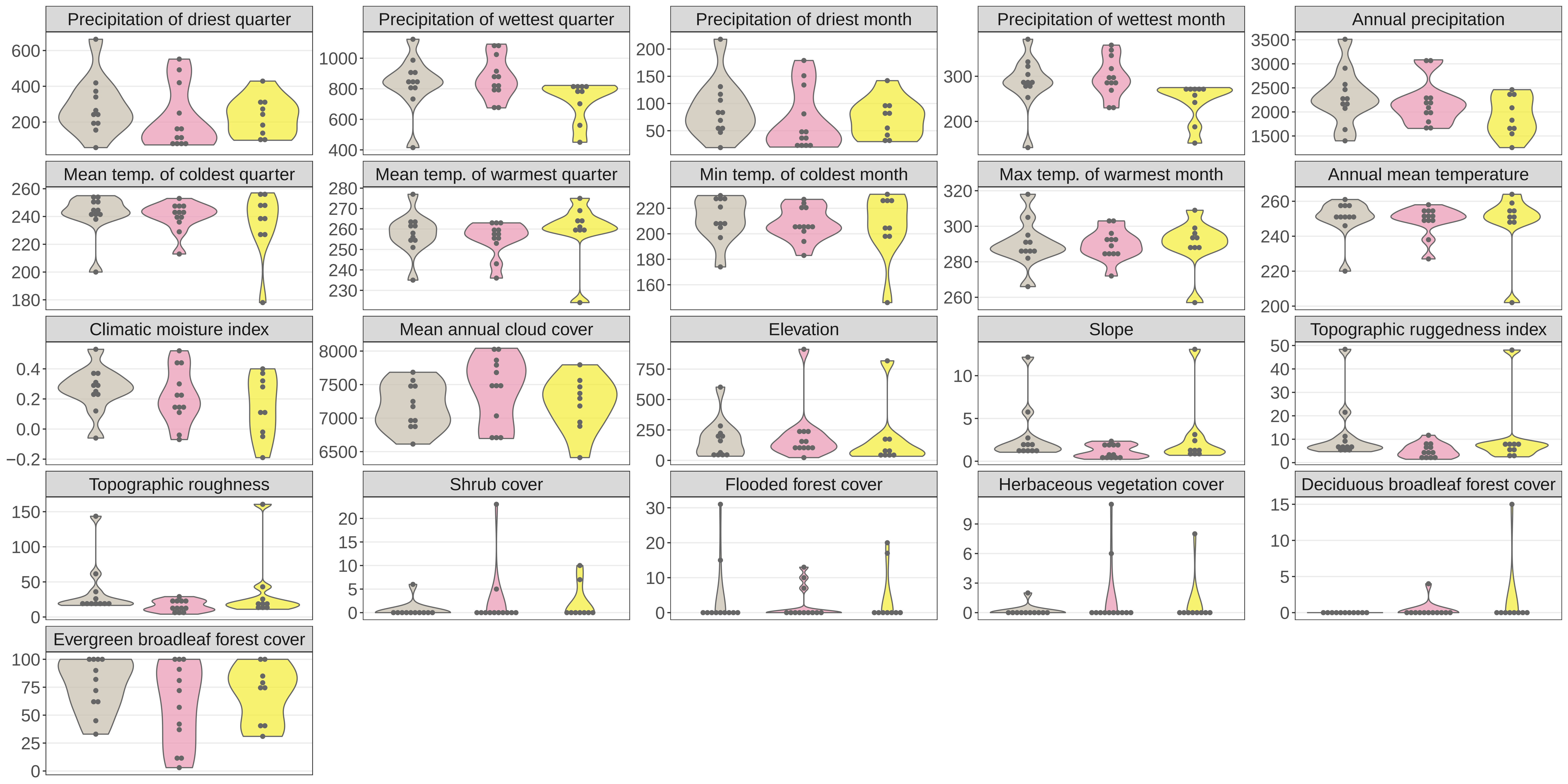
